# *Wolbachia*-induced cytoplasmic incompatibility produces heritable chromatin modifications that suppress position-effect variegation

**DOI:** 10.64898/2026.06.12.731975

**Authors:** Hunter J. Hill, William Sullivan, Brandon S. Cooper

**Author notes:** Corresponding authors: (HJH), (BSC).

## Abstract

Maternally transmitted *Wolbachia* often cause cytoplasmic incompatibility (CI), a sperm modification that kills host embryos lacking the endosymbiont. CI produces defects in paternal chromosome replication, condensation, and segregation during the first zygotic cell cycle, but a significant fraction of embryos progress normally through this and subsequent cycles and only exhibit defects at later developmental stages. These results, together with documented CI-induced epigenetic chromatin modifications, suggest heritable chromatin modifications are responsible for the developmentally delayed defects. Here, we conducted a Position-Effect Variegation (PEV) screen in *Drosophila melanogaster* using *In(1)w^m4^*to test for persistent effects on heterochromatin-mediated silencing in adults that survived CI. We show that *Wolbachia* acts as a variegation suppressor, or *Su(var*), increasing eye pigment when present in CI-inducing fathers, a reproducible effect observed across several maternal genotypes that differed in CI strength. That is, passage of the *In(1)w^m4^*through *Wolbachia*-infected males limits the spread of heterochromatin into the neighboring euchromatin in the progeny. This effect is consistent with disruption of heterochromatin establishment at the mid-blastula transition, when stochastic spreading of heterochromatin determines whether the displaced *white* gene is silenced. Surprisingly, maternal *Wolbachia* did not revert the PEV modification, and in one genotype, *Wolbachia* increased suppression. Together, our results demonstrate that *Wolbachia*-mediated chromatin effects persist to adulthood, are not corrected by CifA-dependent rescue, and can be compounded by maternal *Wolbachia*. These findings establish that rescue is incomplete at the level of heterochromatin-mediated silencing and suggest that CI-specific and constitutive *Wolbachia* chromatin effects may operate through at least partially independent pathways.

## Introduction

Insects often carry maternally inherited *Wolbachia* bacteria that affect their reproduction (Hilgenboecker *et al*., 2008; Hurst *et al*., 1999; Negri *et al*., 2006; Weeks & Breeuwer, 2001; Weinert *et al*., 2015; Werren *et al*., 1995, 2008; Yen & Barr, 1971). The most common reproductive effect – cytoplasmic incompatibility (CI) – kills embryos lacking *Wolbachia* when they are fertilized by *Wolbachia*-modified sperm (Hoffmann *et al*., 1986; Shropshire *et al*., 2020; Wright & Barr, 1981; Yen & Barr, 1973). Rescue occurs when females also carry *Wolbachia* (Hoffmann *et al*., 1986), facilitating spread of the endosymbiont to high frequencies in host populations (Hoffmann *et al*., 1990). CI strength (*i.e.,* the proportion of embryos that die in a CI cross) varies widely among *Wolbachia*-host systems (*e.g.,* Shropshire *et al*., 2022), with relatively weak CI that depends on male age observed for *w*Mel in *Drosophila melanogaste*r (Hoffmann, 1988; Reynolds & Hoffmann, 2002; Shropshire *et al*., 2021). However, *w*Mel causes strong CI in *Aedes aegypti* backgrounds (Walker *et al*., 2011), maintaining the pathogen-blocking transinfection at high frequencies for dengue control (Hoffmann *et al*., 2011; Lenharo, 2023b, 2023a; Utarini *et al*., 2021; Velez, Tanamas, *et al*., 2023; Velez, Uribe, *et al*., 2023). That *w*Mel causes strong CI in novel hosts (*e.g.,* also in *D. simulans;* Poinsot *et al*., 1998), but weaker CI in its native host, supports the evolution of *D. melanogaster*-linked CI suppressors (Turelli, 1994). Relatively weak *w*Mel CI in *D. melanogaster* also provides the opportunity to ask whether CI leaves lasting consequences for the many adults that escape it.

*Wolbachia* are not found in mature sperm (Binnington & Hoffmann, 1989; Snook *et al*., 2000), but CI factors (Cifs) are known to alter paternal chromosomes (Horard *et al*., 2022; Landmann *et al*., 2009). Two *Wolbachia* prophage-associated genes (*cifA* and *cifB*) cause and one gene (*cifA*) rescues CI (Beckmann *et al*., 2017; LePage *et al*., 2017; Shropshire *et al*., 2018). During CI, the CifB nucleomodulin protein interacts with paternal chromosomes, rendering them incompatible with maternal chromosomes in the embryo. Embryonic lethality via CI can occur during the first zygotic mitosis, where the paternal genome is incompletely condensed and DNA replication by metaphase is incomplete (Landmann *et al*., 2009; Tram *et al*., 2003). Anaphase progresses before the paternal chromosomes are resolved, leading to lagging chromosomes (or paternal-chromatin mass), chromosome bridging, and fragmentation (Horard *et al*., 2022; Landmann *et al*., 2009); and CI-derived embryos that are destined to die are often terminated

A number of studies have found that a significant fraction of CI-derived embryos progress normally beyond the initial embryonic divisions and exhibit developmentally delayed post-hatching lethal phases (Bonneau *et al*., 2018; Callaini *et al*., 1996, 1997; Duron & Weill, 2006; Jost, 1970; Lassy & Karr, 1996; Wright & Barr, 1981). This late lethality was attributed to haploidy from first-division chromosome loss, but recent work demonstrated that deferred CI defects are distinct from – and independent of – the CI-induced first-division errors (Warecki *et al*., 2022). Importantly, these delayed defects are eliminated in Rescue crosses (infected females crossed to infected males). For example, approximately one-third of the *D. simulans* embryos derived from a *w*Ri CI cross develop as normal diploids through the blastoderm, undergoing division defects and abnormal development only after cellularization (Warecki *et al*., 2022). Similarly, *D. melanogaster* larvae derived from CI crosses exhibit increased locomotor defects and lethality, with both eliminated in Rescue crosses (Perez *et al*., 2026). These *Wolbachia*-induced delayed defects suggest the involvement of epigenetic mechanisms. This is supported by the observation that the heritable chromatin mark H3K27me1 is elevated in CI-derived embryos (Lee *et al*., 2023; Perez *et al*., 2026).

To determine whether *Wolbachia*-induced CI leaves persistent effects on heterochromatin-mediated silencing in *D. melanogaster*, we assayed gene expression using Position-Effect Variegation (PEV), the stochastic silencing of a gene relocated adjacent to heterochromatin by a chromosomal rearrangement (Spofford, 1968, 1976; Wallrath & Elgin, 1995) (reviewed in Ashburner *et al*., 2005; Eissenberg & Reuter, 2009; Elgin & Reuter, 2013; Weiler & Wakimoto, 1995). We reasoned that epigenetic modifications (Lee *et al*., 2023; Perez *et al*., 2026) affecting the sperm of CI-inducing fathers may be long-lasting and have functional effects in adult-host escapers. To test this, our screen uses *In(1)w^m4^*, with breakpoints in 3B5 and heterochromatin block h28 generating a large inversion that places *white* near pericentric heterochromatin (Muller, 1930; Solodovnikov & Lavrov, 2022). During the mid-blastula transition, each cell undergoes a stochastic heterochromatin spreading—extensive spreading transcriptionally silences the neighboring *white* gene, whereas limited spreading leaves the *white* gene active. Once established, the chromatin state is maintained through development, resulting in clonal white and red patches in the cells of the compound *Drosophila* eye. We reasoned that analysis of these clonal patches in *In(1)w^m4^*-bearing adults derived from CI crosses could provide

To assay PEV in the context of CI, we generated *D. melanogaster* stocks with and without *w*Mel *Wolbachia* that carry *In(1)w^m4^*. We test whether observed epigenetic marks in CI-derived embryos (Lee *et al*., 2023; Perez *et al*., 2026) have persistent effects on heterochromatin-mediated silencing in adults that survive CI. We show that survivors exhibit suppressed variegation of the paternally derived *In(1)w^m4^ X*-chromosome. This demonstrates that during spermatogenesis *Wolbachia* induces heritable chromatin modifications such that some 14 mitotic cycles later during the mid-blastula transition, spreading of the *X*-centric heterochromatin is inhibited, facilitating increased transcription of the neighboring euchromatin genes. We also find this effect persists in Rescue crosses (symbiotic males by symbiotic females) indicating it involves factors distinct from the Cifs (CI and Rescue factors).

## Results

### Outcrosses reveal *Su(var)* mutations in maternal backgrounds

We used light microscopy and an established machine-learning method to assess variegation (**Supp Fig 1**) in the *Drosophila* eye (Hill *et al*., 2025), which is composed of ∼700 ommatidia that import pigment via the *white*-encoded ABC transporter (**Fig 1A–C**). To monitor *Wolbachia-*induced PEV effects on the paternal-*X* chromosome, we designed outcrosses for *In(1)w^m4^* (see **Fig 1D**) males that would generate patroclinous sons (*X* inherited from father) or daughters heterozygous for *white^−^*. Importantly, in both cases, eye pigment comes only from the variegating paternal-*X* chromosome. All fly crosses and development occurred at 26°C (see Materials and Methods).

**Figure 1.**
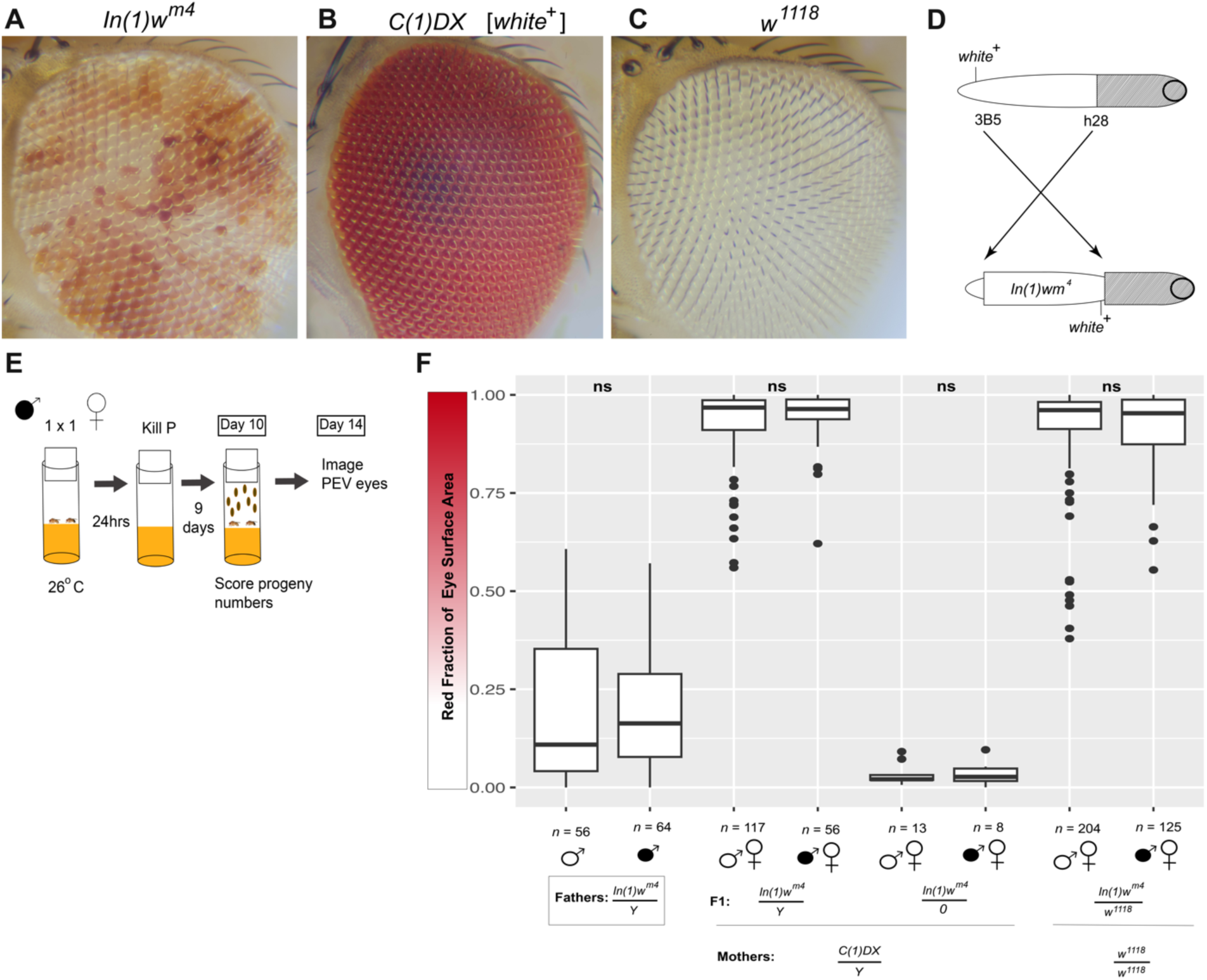
Maternal *Su(var)* mutations cause highly pigmented eyes that preclude analysis of *Wolbachia* effects on the paternal *X*. (**A**) Eye of an *In(1)w^m4^/Y* male, showing variegation. (**B**) Eye of a *C(1)DX /Y* [*white⁺*] female, uniformly pigmented. (**C**) Eye of a *w*^1118^*/Y* male, lacking ocular pigment. (**D**) Diagram of the inversion that generated the *In(1)w^m4^* chromosome: *white* is displaced from its euchromatic position and relocated toward pericentric heterochromatin (grey); circles denote centromeres; breakpoints are labeled at 3B5 and h28. (**E**) Crossing scheme for the pre-introgression PEV screen. Single-pair matings of *In(1)w^m4^/Y* males (± *w*Mel) to *Wolbachia*-free virgin females; progeny scored at day 14. (**F**) Image analysis results from screen plotted as red fraction of eye surface area. Each point is one eye image; sample sizes and genotypes are indicated below. Filled sex symbols denote *Wolbachia*, open symbols denote *Wolbachia*-free. F_1_ progeny showed severely suppressed variegation regardless of paternal *Wolbachia* status, and no cross direction differed significantly (Kruskal-Wallis

Given there are over 500 documented mutations in ∼150 fly genes known to affect PEV (Ebert *et al*., 2004; Schotta *et al*., 2003), we first tested for the presence of *Su(var)* mutations in maternal backgrounds. Young (0-18hrs old) *In(1)w^m4^/Y* males were crossed to *Wolbachia-*free virgin females bearing an attached-*X* chromosome (*C(1)DX*) and *Y* chromosome (*C(1)DX/Y*), and the resulting patroclinous sons were scored (1Mx1F, **Fig 1E**). Most sons were *In(1)w^m4^/Y* (∼86%); however, the remaining sons (∼14%) lacked a *Y* chromosome. *In(1)w^m4^/Y* males could be identified by having significantly more pigment than their fathers, regardless of the father’s *Wolbachia* status. Notably, we could not detect significant differences in variegation between cross directions due to the severe suppression phenotype (**Fig 1F**).

We then outcrossed young *In(1)w^m4^/Y* males to naturally *Wolbachia-*free *w^1118^/w^1118^* only ocular pigment in heterozygous *In(1)w^m4^/w^1118^* females is imported via gene product from the paternally inherited *In(1)w^m4^* chromosome. As with patroclinous sons, heterozygous daughters had severely suppressed variegation compared to their fathers (**Fig 1F**, **Table 1**).

**Table 1.**
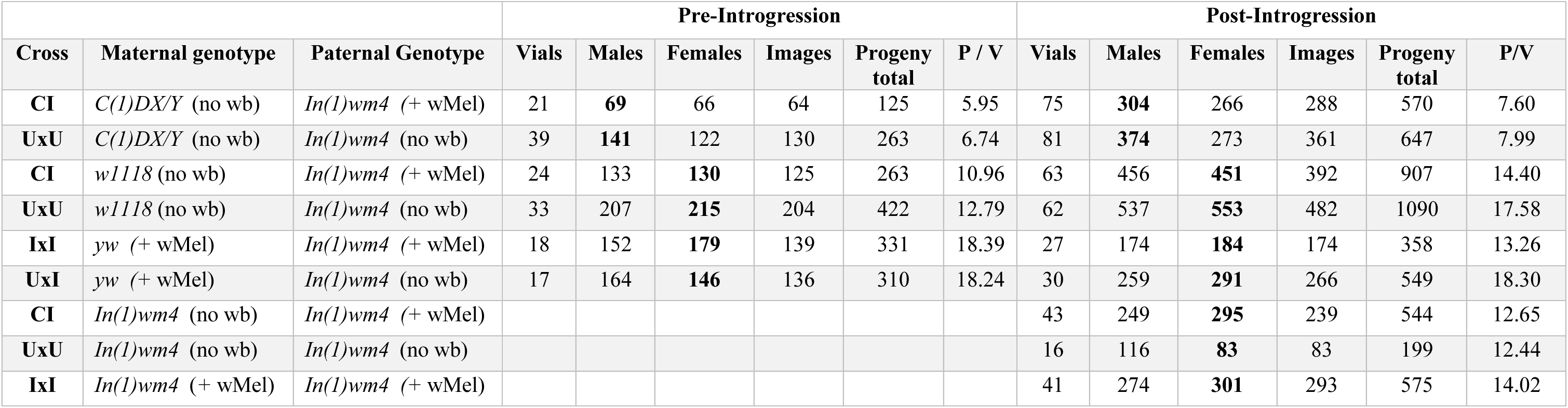
Cross directions, progeny counts, and imaging totals for the PEV screens. Each row gives a parental cross wi genotype and *Wolbachia* status (no wb, *Wolbachia*-free; + *w*Mel, infected). Columns report the number of fertile vials (>0 of males, females, and imaged eyes, total progeny, and mean progeny per vial (P/V), separately for the pre-introgression sc post-introgression screens (**Fig 2D**, **Fig 4**). For each cross, the bolded sex is the F1 class carrying the variegating paternally chromosome and therefore scored for PEV (patroclinous sons for *C(1)DX*; heterozygous daughters for *w^1118^*, and *yw*; and h for *In(1)w^m4^*). Across all scored classes, 90.9% of relevant F1 eyes were imaged (3,376/3,716); unimaged flies died in the m damaged during mounting. Cross abbreviations: CI, *Wolbachia*-infected father × uninfected mother; UxU, uninfected × un infected × infected (Rescue); UxI, uninfected father × infected mother (Reciprocal).

| Cross | Maternal genotype | Paternal Genotype | Pre-Introgression |  |  |  |  |  | Post-Introgression |  |  |  |  |  |
| --- | --- | --- | --- | --- | --- | --- | --- | --- | --- | --- | --- | --- | --- | --- |
|  |  |  | Vials | Males | Females | Images | Progeny total | P / V | Vials | Males | Females | Images | Progeny total | P/V |
| **CI** | *C(1)DX/Y* (no wb) | *In(1)wm4* (+ wMel) | 21 | **69** | 66 | 64 | 125 | 5.95 | 75 | **304** | 266 | 288 | 570 | 7.60 |
| **UxU** | *C(1)DX/Y* (no wb) | *In(1)wm4* (no wb) | 39 | **141** | 122 | 130 | 263 | 6.74 | 81 | **374** | 273 | 361 | 647 | 7.99 |
| **CI** | *w1118* (no wb) | *In(1)wm4* (+ wMel) | 24 | 133 | **130** | 125 | 263 | 10.96 | 63 | 456 | **451** | 392 | 907 | 14.40 |
| **UxU** | *w1118* (no wb) | *In(1)wm4* (no wb) | 33 | 207 | **215** | 204 | 422 | 12.79 | 62 | 537 | **553** | 482 | 1090 | 17.58 |
| **IxI** | *yw* (+ wMel) | *In(1)wm4* (+ wMel) | 18 | 152 | **179** | 139 | 331 | 18.39 | 27 | 174 | **184** | 174 | 358 | 13.26 |
| **UxI** | *yw* (+ wMel) | *In(1)wm4* (no wb) | 17 | 164 | **146** | 136 | 310 | 18.24 | 30 | 259 | **291** | 266 | 549 | 18.30 |
| **CI** | *In(1)wm4* (no wb) | *In(1)wm4* (+ wMel) |  |  |  |  |  |  | 43 | 249 | **295** | 239 | 544 | 12.65 |
| **UxU** | *In(1)wm4* (no wb) | *In(1)wm4* (no wb) |  |  |  |  |  |  | 16 | 116 | **83** | 83 | 199 | 12.44 |
| **IxI** | *In(1)wm4* (+ wMel) | *In(1)wm4* (+ wMel) |  |  |  |  |  |  | 41 | 274 | **301** | 293 | 575 | 14.02 |

These results suggest that both *w^1118^* and *C(1)DX* stocks, carry *Su(var)* mutations. To enable analysis of PEV alterations in CI-derived progeny, we designed a series of crosses to remove *Su(var)s* by genetic introgression of the *Y-*chromosome and the three autosomes.

### Paternal *Wolbachia* suppresses variegation across introgressed maternal backgrounds

We introgressed the *In(1)w^m4^* genetic background into the *w^1118^* and *C(1)DX* stocks (**Fig 2A**) and preserved variegation in the *In(1)w^m4^* stocks through repetitive rounds of selection. Each *In(1)w^m4^* stock (+ or – *w*Mel) was split into two lines: one maintained without selection, and one selectively bred to prevent the accumulation of spontaneous suppressor mutations (**Fig 2B&C**) (S. H. Wang & Elgin, 2019). If less pigmented flies are not selected each generation, the eye color of the population accumulates red pigment, presumably due to fitness deficits in flies with eyes lacking pigment (Ferreiro *et al*., 2018; Rickle *et al*., 2025; Xiao *et al*., 2017).

**Figure 2.**
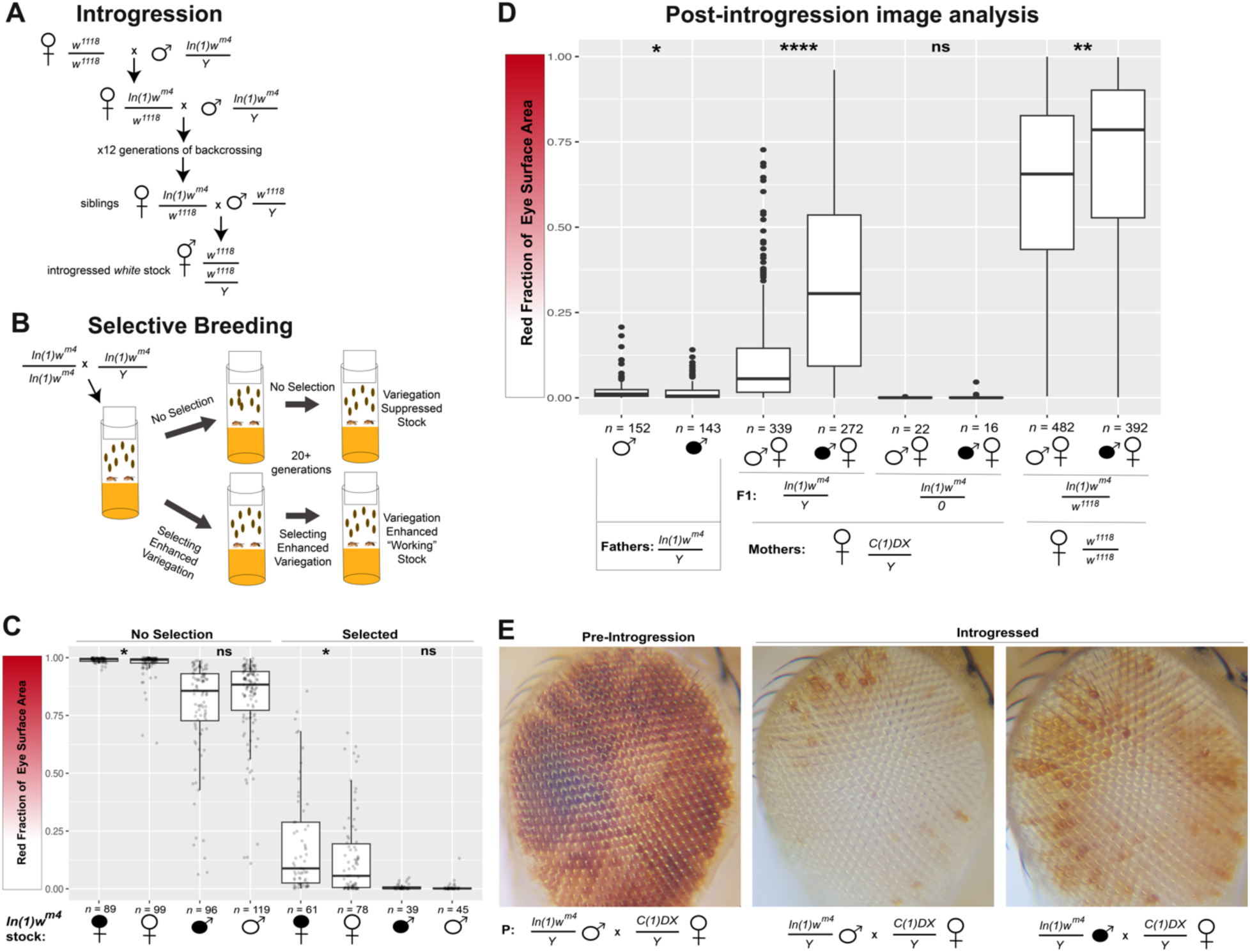
Introgression and selective breeding reveal *Wolbachia*-associated variegation suppression across maternal genotypes. (**A**) Crossing scheme for 12-generation introgression of the *In(1)w^m4^* background into *w*^1118^; *C(1)DX* was introgressed identically (**Supp Fig 2A**). (**B**) Selective breeding scheme used to maintain variegation in *In(1)w^m4^* stocks. (**C**) Eye pigment quantified from images in *In(1)w^m4^* stocks with and without selection. Within both the unselected and selected stocks, one sex comparison was significant (*p*₁ = 0.046; *p*₃ = 0.046) and one was not (*p*₂ = 0.26; *p*₄ = 0.26) (Kruskal-Wallis χ² = 534.78, df = 7, *p* < 2.2 × 10⁻¹⁶; pairwise Mann-Whitney, BH-adjusted). (**D**) Post-introgression PEV screen, red fraction of eye surface area. Left to right: *In(1)w^m4^* fathers differed by *Wolbachia* status ( *p₁* = 0.047)*; C(1)DX* derived (*In(1)w^m4^ /Y*) patroclinous sons showed strongly increased pigment in CI vs. Control *( p*₂ = 2.48 × 10⁻¹⁷); *C(1)DX* derived *In(1)w^m4^/O* males did not differ (ns, *p*₃ = 0.92); *w*^1118^ heterozygous daughters showed increased pigment in CI vs. Control ( *p*₄ = 0.0048) (beta GLMM, BH-adjusted). Sample sizes indicated below each group. (**E**) Representative eye images (near population mean) of F_1_ patroclinous *In(1)w^m4^/Y* males from pre-introgression and introgressed *C(1)DX* crosses. Filled sex symbols denote *Wolbachia*, open symbols denote *Wolbachia*-free.

We anticipated that the *Su(var)* mutations were removed from the genetic backgrounds of *w^1118^,* and *C(1)DX* after introgression. Indeed, when we repeated the PEV screen, the F_1_ variegation was less suppressed in every direction compared to the pre-introgression results (**Fig 1F** compared to **Fig 2D**). Furthermore, after controlling the genetic background, we could also detect a significant variegation effect in progeny derived from *In(1)w^m4^* fathers with *Wolbachia*, compared to those derived from aposymbiotic *In(1)w^m4^* fathers (CI and Control respectively). For *C(1)DX,* the CI cross resulted in 25% increased median pigment compared to the Control, and for *w^1118^*, the CI cross resulted in 13% increased median pigment compared to the Control.

The *C(1)DX* cross (*C(1)DX/Y* x *In(1)w^m4^/Y* ) had the strongest response to the removal of *Su(var)* mutations (**Fig 2E**). The pigment change in F_1_ males was so dramatic that it was no longer possible to differentiate *In(1)w^m4^/Y* and *In(1)w^m4^/O* males visually. To distinguish *Y*-bearing males, we used PCR to amplify *Y*-specific sequences with primers targeting the first exon of *kl-3* (Bernardo Carvalho *et al*., 2000). During eye imaging, whole flies were saved, genomes extracted, and each given a barcode ID to later pair images with *kl-3* PCR results (see Materials and Methods, **Supp Fig 2B, Supp Table 1**). Male genomes that did not successfully amplify the *kl-3* exon were removed from the *In(1)w^m4^/Y* dataset and placed in their own *In(1)w^m4^/O* group (**Fig 2D**). We discovered a significant increase in pigmentation in the eyes of patroclinous sons derived from a CI cross (*In(1)w^m4^/Y* with *Wolbachia* x aposymbiotic *C(1)DX/Y*) compared to the Control (aposymbiotic *In(1)w^m4^/Y* x aposymbiotic *C(1)DX/Y*) (BH-adjusted *p* = 2.48e-17).

The *w^1118^* stock was also purged of *Su(var)* mutations by backcrossing to aposymbiotic *In(1)w^m4^ /Y* males. However, in this PEV screen, CI F_1_ females (heterozygous for *w^1118^/In(1)w^m4^* ) had more pigment (median= 78.5% red) than CI patroclinous *In(1)w^m4^/Y* males (median= 30.8% red). This difference between maternal genotype contributions to PEV pigmentation suggests since most of the *w^1118^* euchromatin was locked in an inversion during introgression. Nonetheless, we again report evidence supporting that there is a moderate, yet significant (*p* = 0.0048), increase in pigmentation in eyes derived from a CI cross (*In(1)w^m4^/Y* with *Wolbachia* x aposymbiotic *w^1118^/w^1118^*) compared to a Control cross (aposymbiotic *In(1)w^m4^/Y* x aposymbiotic *w^1118^/w^1118^*) (**Fig 2D**).

### CI strength does not covary with ocular pigment

We next investigated whether the magnitude of PEV suppression observed in adults that survived CI was correlated with CI strength. To test this, we quantified egg-to-adult viability for a *yw*/*yw* x *In(1)w^m4^/Y* cross prior to imaging the eyes of F_1_ females that survived CI (**Fig 3**, see Materials and Methods). CI crosses showed significantly reduced viability relative to controls (β = 0.451on the log-odds scale, SE = 0.22, *p* = 0.04), corresponding to a 1.57-fold reduction in odds of survival (**Table 2**). Table 2 reports the results of other crosses that showed similar reductions. Vial-level mean eye pigment and egg-to-adult viability were not correlated for CI (r = 0.13, *p* = 0.37; **Figs 3C,S3C**) or Control crosses (r = 0.1, *p* = 0.44 **Figs 3D&S3D**). Thus, we found no evidence that the magnitude of PEV suppression scales with CI strength.

**Figure 3.**
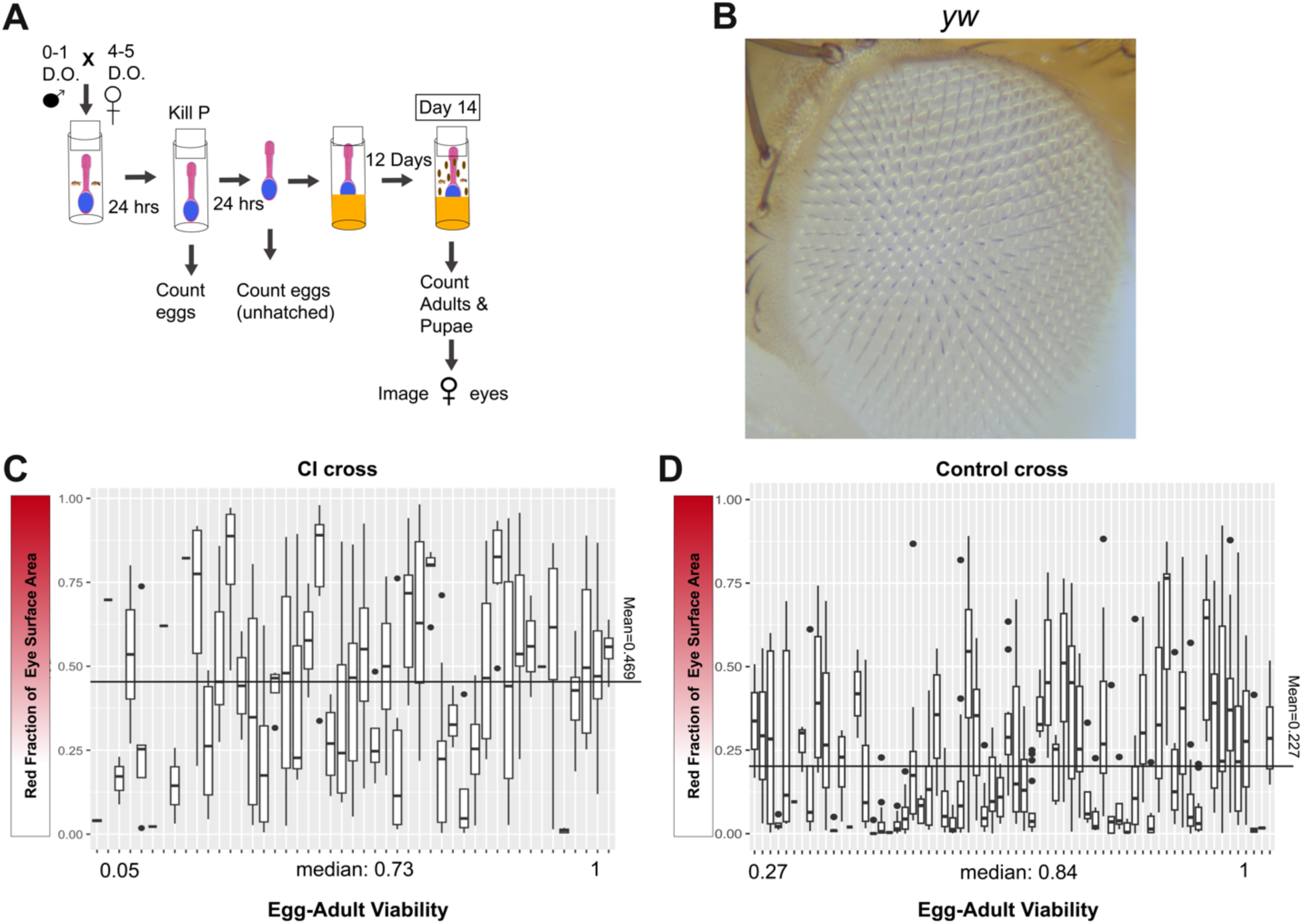
CI strength does not covary with variegation suppression. (**A**) Scheme for the combined CI-strength and PEV assay. Single-pair matings of tetracycline-cured *yw/yw* virgin females to young *In(1)w^m4^/Y* males; eggs counted at 24 h, unhatched embryos counted at 48 h, and adults/pupae scored before imaging F_1_ female eyes at day 14. (**B**) Representative *yw*-cross eye lacking ocular pigment. (**C**) PEV quantification for individual CI-cross vials, each box represents a single vial, ordered along the *x*-axis by egg-to-adult viability (vial median viability 0.73). The horizontal line marks the overall mean red fraction (0.469). (**D**) Same for Control-cross vials (vial median viability 0.84; mean red fraction 0.227). Vial-level mean pigment did not correlate with egg-to-adult viability (Pearson; CI: *r* = 0.13, *p* = 0.37; Control: *r* = 0.10, *p* = 0.44; **Supp Fig 3**). CI crosses showed significantly reduced viability (binomial GEE, β = 0.451, SE = 0.22, *p* = 0.04; **Table 2**) yet significantly higher pigment than Controls (0.469 vs. 0.227, *p* = 7.6 × 10⁻¹⁰).

**Table 2.** Egg-to-adult viability assay quantifying CI strength in three aposymbiotic maternal genotypes. S *In(1)w^m4^/Y* fathers (+ *w*Mel for CI crosses; *Wolbachia*-free for UxU Control crosses) to tetracycline-cured *w^1118^*, mothers. Columns report the number of egg-laying spoons (>5 eggs), total eggs, F_1_ adults, egg-hatch proportion comparison only), and egg-to-adult viability (the CI metric used here, because larval lethality contributes to CI-r *al.*, 2026). Viability was modeled with a binomial generalized estimating equation (logit link, exchangeable corr day as clustering factor); β (log-odds), standard error, odds ratio, and *p* are reported for each CI–Control compar indicate the factor by which Control survival odds exceed CI (e.g., 2.1-fold for *w^1118^*). All three genotypes showe reduced viability in CI crosses (*w^1118^*: *p* = 0.002; *yw*: *p* = 0.04; *In(1)w^m4^*: *p* = 0.016). *C(1)DX/Y* mothers were not strength because ∼half of their embryos are inviable *Y/Y* or *XX/X* genotypes.

| Cross | Maternal genotype (wb-) | Spoons (>5 eggs) | Eggs | F1 Adults | Egg hatch | Egg-Adult Viability | $\beta$ | SE | odds ratio | p value |
| --- | --- | --- | --- | --- | --- | --- | --- | --- | --- | --- |
| CI | *w<sup>1118</sup>* | 41 | 1070 | 700 | 0.808 | 0.654 | 0.741 | 0.24 | 2.1 | 0.002 |
| UxU | *w<sup>1118</sup>* | 51 | 1329 | 1066 | 0.937 | 0.802 |  |  |  |  |
| CI | *yw* | 48 | 855 | 584 | 0.876 | 0.683 | 0.451 | 0.22 | 1.57 | 0.04 |
| UxU | *yw* | 68 | 1353 | 1045 | 0.953 | 0.772 |  |  |  |  |
| CI | *In(1)w<sup>m4</sup>* | 26 | 476 | 285 | 0.811 | 0.599 | 0.597 | 0.25 | 1.82 | 0.016 |
| UxU | *In(1)w<sup>m4</sup>* | 30 | 627 | 461 | 0.915 | 0.735 |  |  |  |  |

As expected, we found significantly more ocular pigment in adults from the CI crosses (*In(1)w^m4^/Y* with *Wolbachia* x aposymbiotic *yw/yw*) compared to the Control crosses (Mean pigment: CI = 0.469, Control = 0.227, *p* = 7.6e-10). Supplemental Figure 4B presents additional data for the *yw* cross prior to introgression.

### Maternal *Wolbachia* does not revert PEV modification and exacerbates it in *yw*

Symbiotic females rescue CI-induced lethality, with indistinguishable chromosome condensation, pronuclear replication, and egg-hatch phenotypes observed for Control, Rescue, and Reciprocal crosses (Landmann *et al*., 2009; Shropshire *et al*., 2021; Warecki *et al*., 2022). Thus, we initially predicted that in our screen that the Control, Rescue, and Reciprocal crosses might have indistinguishable ocular pigment. When we repeated the screen with symbiotic *yw/yw* females and examined the combined image data we observed an additive *Wolbachia* effect. Control crosses had the least amount of pigment, while Rescue crosses had the most (**Fig 4A**, median values: Control= 0.137; CI= 0.466; Reciprocal= 0.529; Rescue= 0.782).

**Figure 4.**
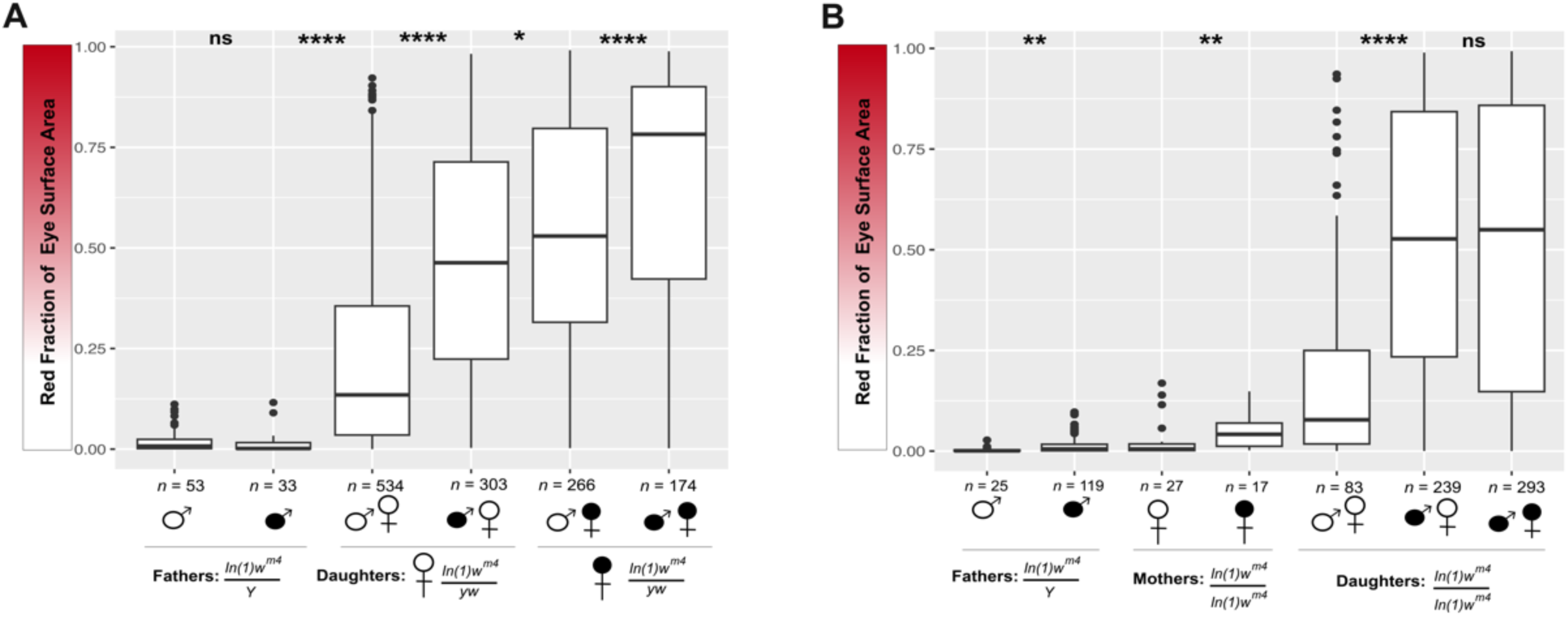
Maternal *Wolbachia* does not rescue the CI-induced PEV phenotype and amplifies it in *yw*. Red fraction of eye surface area; each point is one eye image, sample sizes indicated below each box. Filled sex symbols denote *Wolbachia*-carrying parents, open symbols denote *Wolbachia*-free; paired symbols indicate paternal then maternal genotype. (**A**) The *yw* experiment. *In(1)w^m4^/Y* fathers’ eye pigmentation did not differ by *Wolbachia* status (open vs. filled ♂; ns, *p*₁ = 0.163). Among F_1_ *In(1)w^m4^/yw* daughters, pigment increased stepwise across the four cross directions (median values: Control = 0.137; CI = 0.466; Reciprocal = 0.529; Rescue = 0.782), with Control significantly exceeding fathers (*p*₂ = 1.53 × 10⁻¹⁴), CI significantly exceeding Control (*p*₃ = 7.58 × 10⁻¹⁰), Reciprocal exceeding CI (*p*₄ = 0.017), and Rescue exceeding Reciprocal (*p*₅ = 6.14 × 10⁻⁵) (Kruskal-Wallis χ² = 500.69, df = 5, *p* < 2.2 × 10⁻¹⁶; pairwise tests BH-adjusted, see Methods). (**B**) The homozygous *In(1)w^m4^/In(1)w^m4^* experiment. *In(1)w^m4^/Y* fathers differed by *Wolbachia* status (*p*₁ = 0.0028) as did *In(1)w^m4^/In(1)w^m4^* mothers (*p*₂ = 0.0039). Among F_1_ *In(1)w^m4^/In(1)w^m4^* daughters, the CI cross showed significantly increased pigment over Control (*p*₃ = 1.31 × 10⁻⁵), but the Rescue cross did not differ from CI (ns, *p*₄ = 0.79), indicating non-rescue without additivity (median values: Control = 0.078; CI = 0.527; Rescue = 0.557) (Kruskal-Wallis χ² = 397.92, df = 6, *p* < 2.2 × 10⁻¹⁶; BH-adjusted).

This observation led us to perform our analysis with *In(1)w^m4^/In(1)w^m4^* females in CI, Control, and Rescue crosses to further characterize the effect of maternally contributed *Wolbachia*. Once again, the F_1_ daughters in a CI cross had significantly increased pigment when compared to the Control cross (**Fig 4B**, median values: Control= 0.078; CI= 0.527; Rescue= 0.557). However, in contrast to *yw,* the Rescue cross did not show evidence of an additive *Wolbachia* effect, nor did it rescue (or revert) the PEV phenotype to the Control level. We conclude that maternally contributed *Wolbachia* does not rescue the PEV modification imparted

### *Wolbachia* alters chromatin silencing without disrupting polytene structure

Our discovery that *Wolbachia* suppresses PEV led us to ask whether the polytene chromosome structure of symbiotic stocks is disrupted by *Wolbachia*. Genetic *Su(var)* mutations can substantially perturb polytene chromosome structure, with defects ranging from misaligned chromatin fibrils to amorphous, clumped morphology (Lu *et al*., 2009). *Wolbachia* appeared polarized in the cytoplasm when we imaged larval salivary glands (**Fig 5A**). After confirming the breakpoints of the *In(1)w^m4^*chromosome in males (**Fig 5B**, see Materials and Methods), we imaged males and females with and without *Wolbachia* focusing on the distal part of the *X-*chromosome since it is easily identified cytologically and measured (**Fig 5C**). Occasionally, the entire *X-*chromosome was separated from the rest of the nucleus by squashing pressure and took on a striking conformation that resembled an inversion loop (Reuter *et al*., 1985; Spofford, 1976). This configuration occurred in males regardless of *w*Mel status, and the inversion evidence for disrupted polytene structures in the distal portion of the *X* where we focused our study, nor did we notice any unnatural puffing or constrictions in other regions of the *X-*chromosome or autosomes that would indicate problems with polytenization caused by *Wolbachia* (**Fig 5E**). This included analysis of polytene chromosomes of offspring from multiple CI crosses (*C(1)DX* × *In(1)w^m4^* and *In(1)w^m4^*× *In(1)w^m4^*), confirming that CI does not affect large-scale polytenization. We also assessed males from a *D. simulans* stock that carries *w*Ri, which causes very strong CI (Hoffmann *et al*., 1986; Shropshire *et al*., 2022; Turelli & Hoffmann, 1995). The *X* polytenization of *D. simulans* also appeared unaffected by *Wolbachia* (**Fig 5E**). We conclude that *Wolbachia* does not have obvious effects on polytene chromosome structure in the focal genotypes assayed here.

**Figure 5.**
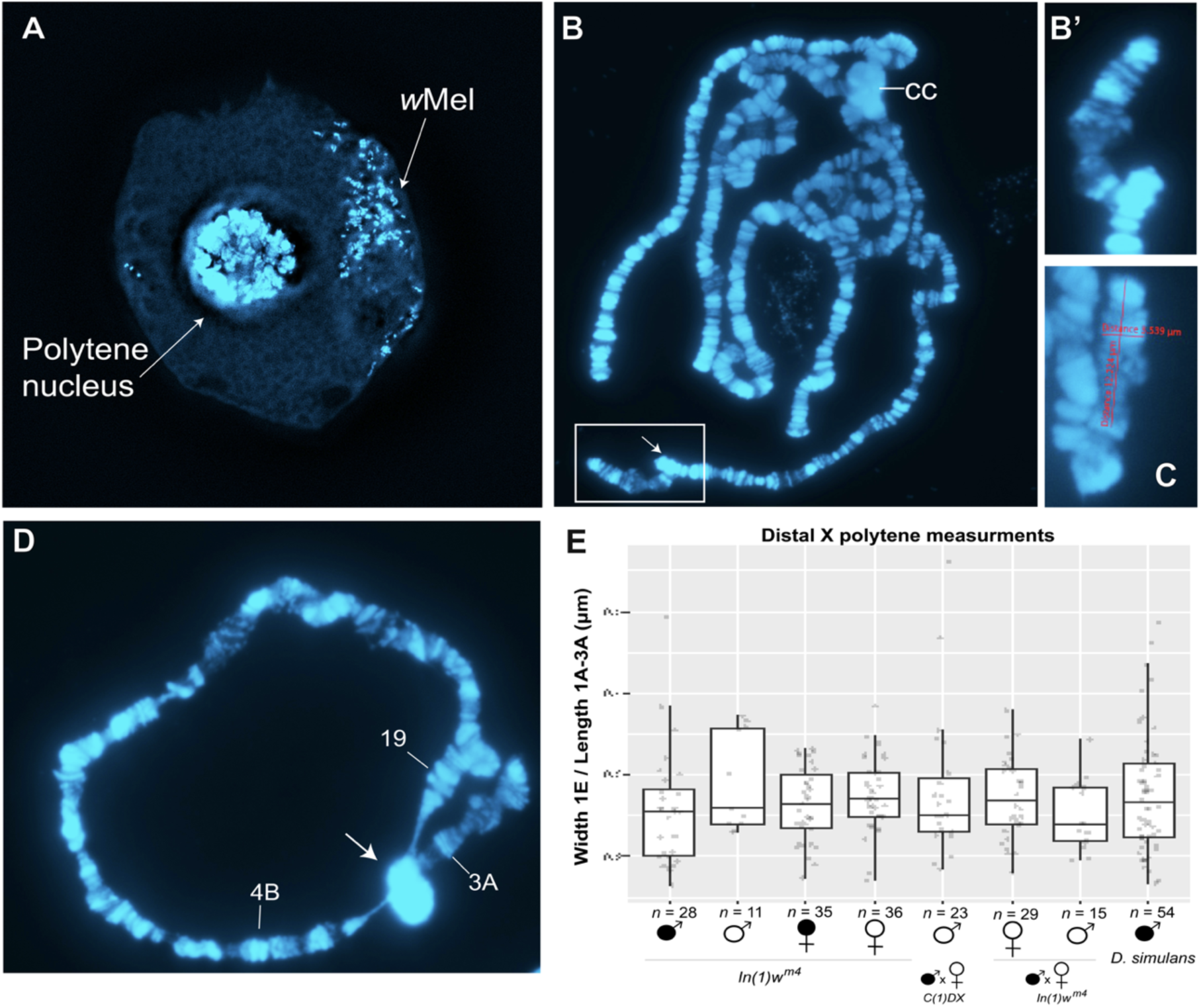
Polytene chromosome morphology is unaffected by *Wolbachia* infection and CI. (**A**) Intact salivary gland secretory cell from a *w*Mel-symbiotic *In(1)w^m4^* larva stained with DAPI; *Wolbachia* (bright cytoplasmic puncta) and the polytene nucleus are marked. (**B**) Squashed polytene nucleus of a *w*Mel-symbiotic

## Discussion

In addition to the well-documented embryonic defects, a number of studies have demonstrated that CI produces distinct developmental defects during the larva-to-adult stages. While these were thought to be delayed defects arising from abnormalities in the first divisions, such as haploid escapers, the CI-induced larva-to-adult phenotypes are now appreciated as distinct and independent from the initial CI-induced embryonic defects. For example, CI-derived *D. simulans* embryos that had progressed normally through the syncytial divisions would exhibit increased segregation errors during the gastrulation mitotic divisions (Warecki *et al*., 2022). In *D. melanogaster*, CI-derived progeny exhibit developmentally delayed larval and adult locomotor defects (Perez *et al*., 2026). That *Wolbachia* effects on sperm produce such delayed defects suggests epigenetic mechanisms are involved. Support for this comes from the finding that *Wolbachia* upregulates H3K27me1 marks on the chromatin of CI-derived embryos (Lee *et al*., 2023; Perez *et al*., 2026). However, those studies could not determine the phenotypic consequences of the CI-induced alterations in epigenetic marks, and it remained unclear if the increased H3K27me1 was responsible for the delayed CI-induced phenotypes.

Thus, this work set out to address the following: Does CI result in altered chromatin states that are inherited through successive mitotic divisions? Do these CI-altered chromatin states have phenotypic consequences? As described, the inverted chromosome *In(1)w^m4^* provides a direct readout of mitotically inherited spreading of the centric heterochromatin (Eissenberg, 1989; Prokofyeva-Belgovskaya, 1947). During gastrulation, the extent of spreading is set by a self-propagating HP1a molecular loop that establishes and maintains the chromatin state into adulthood (Yuan & O’Farrell, 2016). Clonal patches of white ommatidia in *In(1)w^m4^* indicate extensive spreading into neighboring euchromatin and transcriptional silencing of the *white* gene (Spofford, 1976; Tartof *et al*., 1984).

After controlling the genetic backgrounds of *In(1)w^m4^*, *C(1)DX*, *w^1118^*, and *yw,* we observed moderate, yet significant variegation suppression in the CI cross relative to Control. When compared to the suppression by background *Su(var)* mutations we uncovered, or the variegation enhancement in *X/O* males, the CI effect is modest. Yet, our ability to recapitulate the results in 4 separate experiments with unique maternal *X*-chromosome genotypes confirms that it is reproducible. Interestingly, within the *yw* genotype, where CI strength was measured per also had the weakest measurable CI of the three genotypes tested (**Table 2**) yet produced the strongest PEV effect. This dissociation suggests that the chromatin-level effect of paternal *Wolbachia* is at least partially independent of the CI pathway leading to embryonic lethality. A similar independence between CI strength and epigenetic state was reported for testes DNA methylation across *Wolbachia* strains (LePage *et al*., 2014).

Although the PEV effect does not covary with CI strength, we find the action of *Wolbachia* during spermatogenesis suppresses spread of the centric *X* heterochromatin into the neighboring euchromatin. Temperature shift PEV studies demonstrate that the heterochromatic state is established and fixed during the late syncytial divisions and cellularization (Hartmann-Goldstein, 1967). This timepoint coincides with the slowing of the cell cycle, genome-wide activation of zygotic transcription, and establishment of facultative heterochromatin (Farrell & O’Farrell, 2014). Previous studies demonstrated that CI-derived embryos exhibited increased failures in sister chromosome segregation during late syncytial divisions and the divisions immediately following cellularization (Warecki *et al*., 2022). Thus, the deferred segregation phenotype and the PEV defect may share an underlying cause in disrupted heterochromatin establishment, as the state of centric heterochromatin plays a key role in the final stages of sister chromosome segregation (Oliveira *et al*., 2014). Taken together, these studies and the work presented here strongly support that, in addition to the documented first-division defects, CI produces a distinct set of defects at cellularization. Significantly, the work presented here clearly connects CI action at this time with a phenotypic consequence: transcriptional inactivation of the *white* gene.

CI-induced PEV suppression also provides clues to the host mechanisms being targeted. PEV-induced heterochromatin spreading is initiated by the methyltransferase SU(VAR)3-9, which methylates H3K9, resulting in the recruitment of HP1a, which in turn recruits more SU(VAR)3-9 methyltransferase onto neighboring histones (Schotta *et al*., 2002). This feedback loop explains heterochromatin spread and establishment. Classic *Su(var)* screens reveal these mutations fall into two classes: the histone-associated proteins such as HP1a, and the enzymatic components such as the methyltransferases, demethylases, and deacetylases (Elgin & Reuter, 2013). Thus, it is likely that CI disrupts the action of either the structural or enzymatic components of heterochromatin formation.

A specific molecular candidate emerges from the mechanism underlying PEV establishment. In early *Drosophila* embryos, Eggless/SetDB1 deposits H3K9me3 on satellite repeats during the progressively lengthening interphases of nuclear cycles 12–14 (Seller *et al*., 2019)—precisely when deferred CI defects arise (Warecki *et al*., 2022). SetDB1 enzymatic activity requires monoubiquitination (Osumi *et al*., 2019; Seller *et al*., 2019), and Type 1 CifB (*i.e.,* CidB) possesses deubiquitylase activity (Beckmann *et al*., 2017, 2021). While a direct CifB-SetDB1 interaction has not been demonstrated, CidB-mediated deubiquitination of SetDB1 could in principle impair H3K9me3 deposition at the mid-blastula transition, reducing heterochromatin formation at the locus where *In(1)wm4* places *white* adjacent to pericentric heterochromatin. *Wolbachia* infection upregulates *egg* (*dSetDB1*) and downregulates *Su(var)3-9* in *Drosophila* ovaries (Kagemann *et al*., 2024). Whether these expression changes extend to the male germline or the early embryo is untested, but the direction of *Su(var)3-9* downregulation is consistent with the PEV suppression we observe.

Additional insight into the mechanism by which PEV is suppressed in CI-derived embryos comes from recent work demonstrating increased levels of H3K27 monomethylation in CI-derived embryos (Lee *et al*., 2023; Perez *et al*., 2026). Increased H3K27me1 levels could contribute to PEV suppression through a number of mechanisms. For example, since H3K27 and H3K9 are part of the same histone tail (Strahl & David Allis, 2000), increased levels of the former may sterically inhibit HP1a recruitment and SU(VAR)3-9 action. Alternatively, H3K27me1 is enriched at actively transcribed loci (L. Wang *et al*., 2018) and may counter the silencing of HP1a/SU(VAR)3-9 complex at the *white* locus. CifB-mediated disruption of SetDB1 activity and H3K27me1-mediated interference with the HP1a/SU(VAR)3–9 feedback loop are not mutually exclusive and could act in concert to suppress heterochromatin spreading at the *In(1)w^m4^*breakpoint. The dynamic nature of PEV, in which heterochromatin undergoes repeated rounds of silencing and de-repression throughout development (Bughio & Maggert, 2022), means that even subtle shifts in the balance of H3K9 methylation or HP1a recruitment could compound over successive cell divisions to produce the reproducible *Su(var)* phenotype we observe. This proposal rests on three converging lines of evidence: (1) the demonstrated association between paternal *Wolbachia* and PEV suppression across multiple maternal genotypes (this study), (2) the documented increase in H3K27me1 on CI paternal chromatin (Lee downregulation of histone modifiers observed in *w*Mel-infected *Drosophila* cell lines (Jacobs *et al*., 2025). Consistent with effects operating at the level of histone modifications rather than large-scale chromatin architecture, we found no evidence that CI grossly affects polytene chromosome structure, including in CI-cross offspring, but future high-resolution experiments focused at the euchromatin-heterochromatin boundaries may reveal subtle differences in compaction overlooked in this analysis (**Fig 5E**).

The CI-induced PEV suppression is distinct from the other documented CI phenotypes in that it is not reversed in the Rescue cross when both parents have *Wolbachia*. In Rescue crosses, maternal CifA rescues embryonic lethality by preventing the acute chromosome segregation defects caused by paternal CifB (Shropshire *et al*., 2018). However, CifA rescue in the embryo may not correct the upstream chromatin modification imprinted during spermatogenesis; the histone retention, protamine deficiency, and DNA damage in sperm are pre-fertilization events that CifA in the egg cannot undo (Kaur *et al*., 2022; Kaur, Mcgarry, *et al*., 2024). PEV suppression in Rescue crosses therefore represents a persistent effect on heterochromatin-mediated silencing that survives even when CifA prevents lethality.

This result bears directly on the two competing mechanistic models for CI. Under the host-modification model, CifB alters sperm chromatin prior to fertilization (Poinsot *et al*., 2003; Shropshire & Bordenstein, 2019); our observation that these effects persist despite CifA-mediated rescue is a natural prediction of this model, since rescue acts in the embryo and cannot retroactively correct a pre-fertilization modification. Under the toxin-antidote (TA) model, CifB is delivered to the embryo by unaffected sperm and its toxicity is neutralized by CifA (Beckmann *et al*., 2019; Horard *et al*., 2022). The TA model predicts that rescue should be comprehensive, which our PEV data contradict at the level of heterochromatin-mediated silencing. Importantly, our results do not resolve whether CifB acts in the testis or the embryo for the purpose of inducing lethality. However, deferred CI defects are distinct from and independent of first-division segregation errors (Warecki *et al*., 2022), consistent with the possibility that CI operates through both toxin-antidote and host-modification mechanisms at different developmental stages.

In *yw* Rescue crosses, where both parents carry *Wolbachia*, PEV suppression was not restored to Control levels but instead exceeded that of the CI cross, producing a stepwise *yw* raises the additional possibility that *Wolbachia* present in the developing female modifies chromatin through a CifA-independent pathway. This interpretation is supported by evidence that *w*Mel infection reduces chromatin contacts and downregulates histone modifier genes independent of CI (Jacobs *et al*., 2025), and by the downregulation of *Su(var)3-9* in *Wolbachia*-infected ovaries independent of CI (Kagemann *et al*., 2024). If *Wolbachia* constitutively loosens host chromatin through these CI-independent mechanisms, the additive pattern we observe in *yw* Rescue crosses would reflect the convergence of two distinct pathways: a paternal CI-specific effect from spermatogenesis, compounded by a maternal constitutive chromatin effect from *Wolbachia* in the developing female. However, the cell culture observations (Jacobs *et al*., 2025) have not been confirmed *in vivo*, and the ovarian expression changes (Kagemann *et al*., 2024) have not been tested in the male germline or early embryo. Distinguishing between these explanations will require testing additional symbiotic maternal genotypes and directly assaying chromatin state on paternal versus maternal chromosomes in CI escapers.

Our results demonstrate that *Wolbachia*-mediated modification of paternal chromatin during spermatogenesis leaves a persistent signature on heterochromatin-mediated silencing that does not covary with CI strength. Maternal *Wolbachia*, which fully rescues CI-induced lethality, does not restore heterochromatin-mediated silencing to Control levels. In the *yw* background, maternal *Wolbachia* compounds rather than corrects the paternal effect, establishing that CI rescue is incomplete at the level of heterochromatin and suggesting that CI-specific and constitutive *Wolbachia* chromatin pathways may converge in doubly exposed offspring.

Together with prior work, our results define the consequences of CI and Rescue across host life stages (**Fig 6**). This work characterizes persistent chromatin effects in *w*Mel’s natural host that are potentially relevant for hosts carrying *w*Mel-like variants (Shropshire *et al*., 2026) and *Ae. aegypti*, which exhibits CI-associated sperm-chromatin disruption when transinfected with *w*Mel (Kaur, Meier, *et al*., 2024). Theoretical models (*e.g.,* Hoffmann *et al*., 1990) and empirical analyses (*e.g.,* Cooper *et al*., 2017; Hague *et al*., 2020; Kriesner *et al*., 2013, 2016) focused on *Wolbachia* population biology have not typically considered post-embryonic effects of CI on host fitness (*e.g.,* Perez *et al*., 2026; Warecki *et al*., 2022). Accounting for such effects has the potential to better predict *Wolbachia* frequencies (Ravikanthachari *et al*., 2026), including in transinfected *Ae. aegypti* populations where strong CI tends to maintain the pathogen-blocking *w*Mel transinfection at high frequencies despite its negative fecundity effects (Hoffmann *et al*., 2011; Utarini *et al*., 2021; Velez, Tanamas, *et al*., 2023; Velez, Uribe, *et al*., 2023).

**Figure 6.**
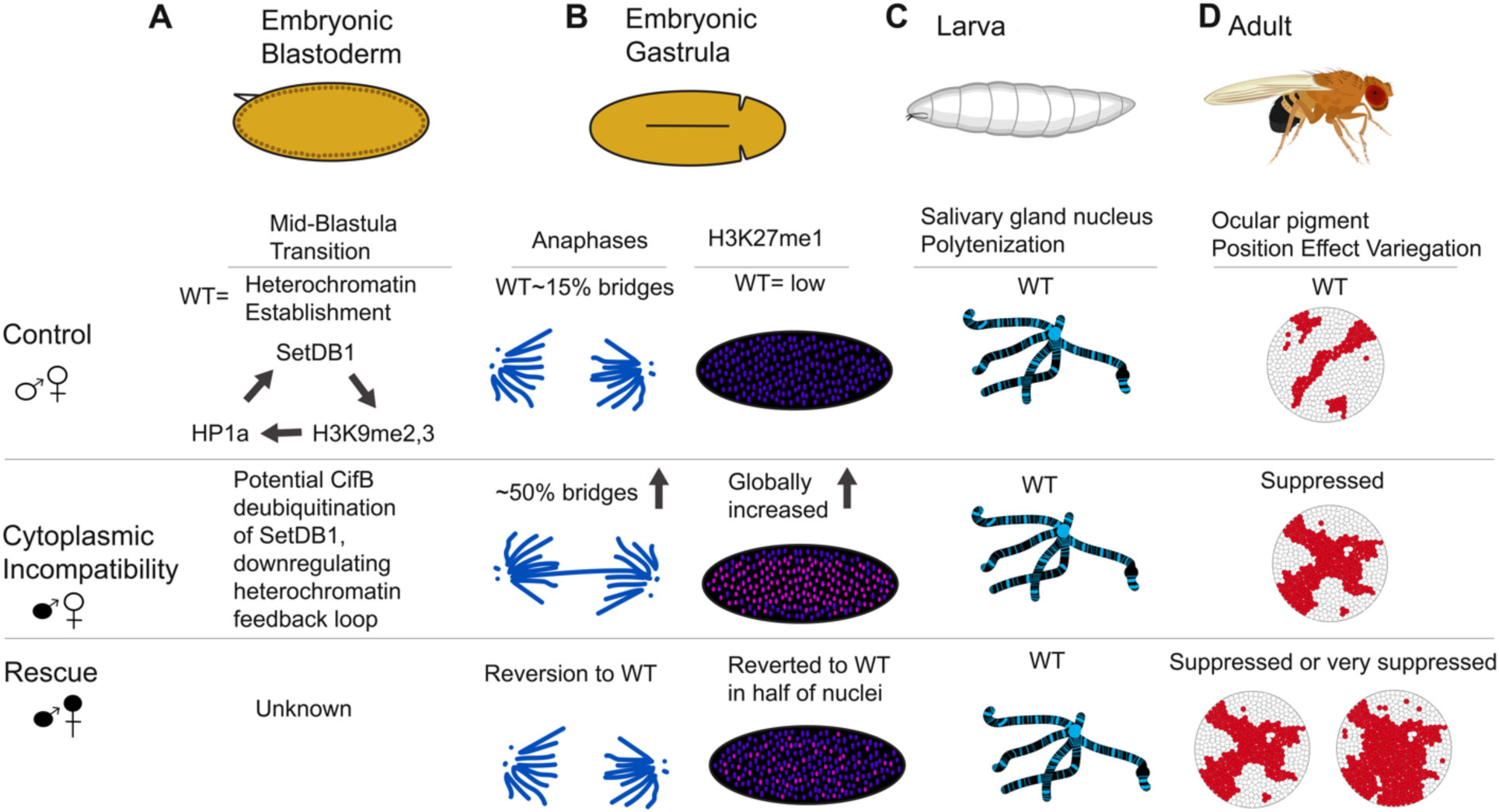
Working model for CI and Rescue across the *Drosophila melanogaster* life cycle. Columns show four life stages (**A** Embryonic Blastoderm, **B** Embryonic Gastrula, **C** Larva, **D** Adult); rows show Control, Cytoplasmic Incompatibility, and Rescue cross directions. (**A**) In the embryonic blastoderm, heterochromatin is normally established through a feedback loop during the mid-blastula transition (division cycles 13-14) which may be disrupted during CI (**B**) Later in embryogenesis, anaphase bridges increase from ∼15% (Control) to ∼50% (CI) and revert toward wild-type in Rescue; H3K27me1 is low in Control, globally increased in CI, and reverted in roughly half of nuclei in Rescue. (**C**) In larvae, salivary gland polytenization is wild-type across all cross directions despite high cytoplasmic *Wolbachia*; CI-derived larvae also show locomotor defects (reverted in Rescue). (**D**) In adults, ocular PEV is wild-type in Control, suppressed in CI, and either not reverted in Rescue (*In(1)w^m4^/In(1)w^m4^*) or further suppressed (*yw/In(1)w^m4^*).

## Materials and methods

### Fly rearing and genotypes

All *Drosophila* stocks were reared at room temperature (∼23°C) on a standard cornmeal-based medium containing yellow cornmeal, dry corn syrup solids, malt extract, inactive dry yeast, soy flour, and agar, with tegosept and propionic acid added as mold and bacterial inhibitors (protocol:Wheeler *et al*., 2024). We discarded flies after 3 weeks since *In(1)w^m4^* stocks aged more than 3 weeks have reduced ocular pigment due to malnutrition. Males and females were retrieved from stocks between 2-3 weeks old. Young males were isolated (0-18hrs old) and crossed to female virgins aged 4-5 days in pairwise matings by aspiration. All fly crosses and development occurred strictly at 26°C since PEV silencing (Gowen & Gay, 1934; Spofford, 1976), *Wolbachia* titer (Hague *et al*., 2022), and CI strength (Bagchi *et al*., 2026; Reynolds *et al*., 2003) can be sensitive to temperature. Mating pairs were removed after 24 hours to ensure mated males remained young.

We introgressed the *In(1)w^m4^* genetic background into the *w^1118^* , *yw*, and *C(1)DX* stocks (**Fig 2A**). In 12 generations, most of the genome was replaced, including the *Y*, where polymorphisms significantly affect heterochromatin and gene expression (Brown *et al*., 2020; Lemos *et al*., 2008, 2010; Paredes *et al*., 2011; Paredes & Maggert, 2009). Notably, during *yw* and *w^1118^* introgression, *In(1)w^m4^* was maintained as a heterozygous inversion which does not form recoverable single-crossover products and thus may retain suppressors (**Fig 2A**). During *C(1)DX* introgression, we replaced the original *FM6* balancer *X-*chromosome with *In(1)w^m4^*(**Supp Fig 2A**).

Maternal genotypes used for PEV screen were *C(1)DX/Y, w^1118^/w^1118^,* and *y^1^ w^1118^/y^1^ w^1118^* (**Table 3**). *C(1)DX/Y* females generate a relatively high proportion of *X/O* male progeny. *In(1)w^m4^/O* males arise from meiotic nondisjunction in the attached-*X* mother resulting in a nullo-*X* egg fertilized by *X-*bearing sperm. PEV males lacking a *Y* chromosome have very little ocular pigment and variegation is enhanced (*i.e.,* silencing increased) (Delanoue *et al*., 2023; Francisco & Lemos, 2014; Gowen & Gay, 1934). Males can be verified as *In(1)w^m4^/O* by their sterility, stellate crystals in the testis (Bozzetri *et al*., 1995), PCR targeting *Y-*specific sequences (Bernardo Carvalho *et al*., 2000), and absence of their *Y* chromosome in the larval karyotype (**Supp Fig 2**). Prior to selective breeding, *In(1)w^m4^/Y* males could be identified by having significantly more pigment than their fathers, regardless of the father’s *Wolbachia* status (**Fig 1F**). Symbiotic *D. melanogaster* stocks were verified to be *w*Mel (as opposed to *w*MelCS, see Materials and methods section below *Wolbachia* status in *Drosophila* stocks)

**Table 3.**
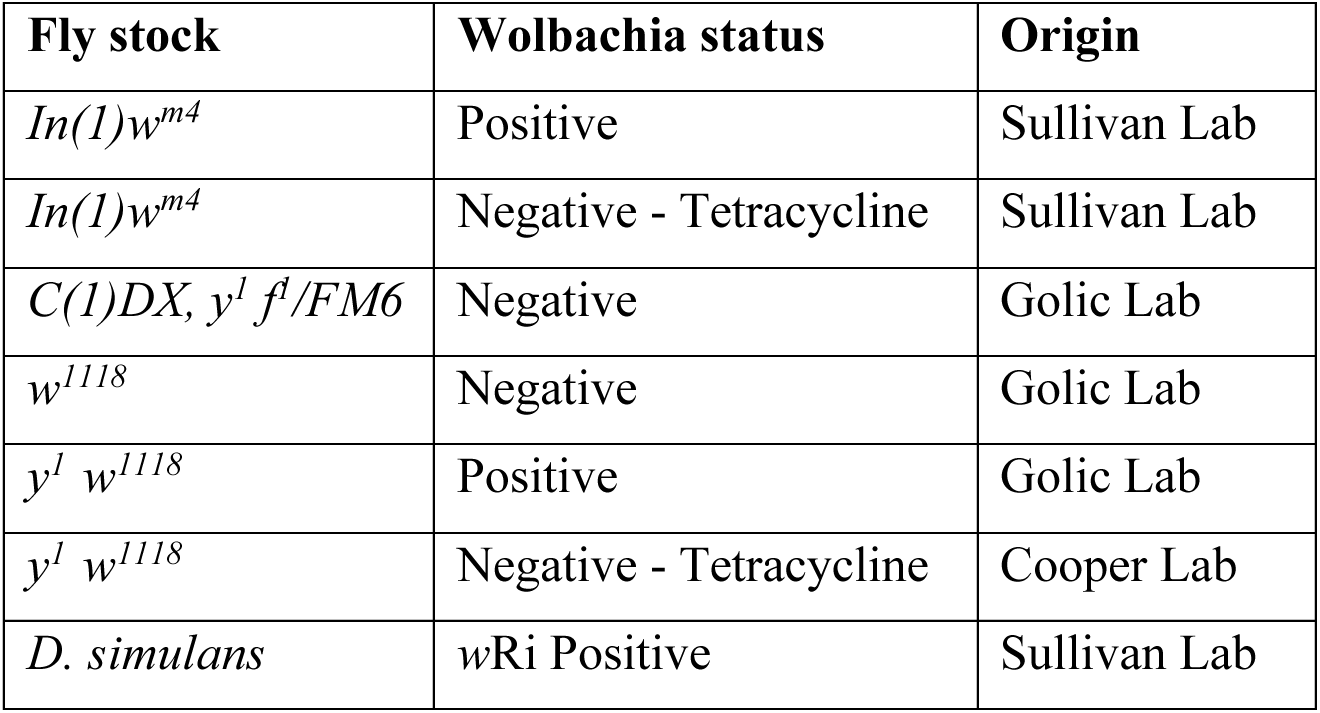
*Drosophila melanogaster* stocks used in this study. For each stock, *Wolbachia* status (Positive; Negat Tetracycline, indicating tetracycline curing) and laboratory of origin are given. *In(1)w^m4^* stocks were reared, *Wol* tetracycline-treated in the Sullivan Lab prior to use in this study. The *C(1)DX*, *w^1118^*, and *yw* stocks were provide *Wolbachia* status was repeatedly verified here, and the *yw* line was tetracycline-cured in the Cooper Lab. Stocks maintained; the *C(1)DX, y^1^ f^1^ / FM6* source stock had its *FM6* balancer replaced by *In(1)w^m4^* during introgressio (see **Fig 2A**, **Supp Fig 2A**).

### Egg-to-adult viability assay

A young (0-1 day old) male and an aged (4-6 days old) female virgin were aspirated into an empty vial along with an ice cream spoon filled with blue fly food and allowed to mate for 24 hours at 26°C. At 24 hours, both parents were removed and euthanized, then embryos were counted. Spoons with <5 embryos were discarded at this stage, and spoons with >5 embryos were returned to the vial for another 24 hours at 26°C. At the 48-hour timepoint, unhatched 26°C. Flies developed over the next 12 days at 26°C, based on our previous crosses, most adults eclose on day 10. 14 days after the start of the experiment, F_1_ flies were counted, and females were immediately imaged.

Pair matings (1×1) between tetracycline-cleared *yw*/*yw* females and *In(1)w^m4^/Y* were performed, and egg-adult viability for each vial was assayed (**Table 2**). The CI cross resulted in 47 fertile vials (with >5 embryos per spoon) and ranged from *N* = 1 to *N* = 20 females per vial with an average of 6.4 females. The Control cross resulted in 67 fertile vials ranging from *N* = 1 to *N* = 26 females per vial with an average of 8.1 females plotted chronologically in **Supp Fig 3A&B**). Although egg hatch rate is typically used to quantify CI, we opted for egg-adult viability as a metric for CI strength, since larval lethality has been shown to contribute to CI-related death in *D. melanogaster* (Perez *et al*., 2026).

### Mottled-eye photography

Four days post-eclosion, adult flies were halved on the anterior-posterior axis and affixed to a standard microscope slide by the thorax. Eyes were imaged on an Olympus BH2 upright brightfield microscope with the 10x objective and a 42 Megapixel mirrorless full-frame Sony DSLR camera. Multiple images (5-9) were captured in the z-plane and stacked together using the program HeliconFocus. Pseudo-3d (stacked) eye photos were reduced from ∼340MB tifs to ∼6MB jpgs in Adobe Lightroom.

### Mottled-eye image quantification

We recently demonstrated that machine learning and light microscopy reliably quantify the PEV phenotype (Hill *et al*., 2025). This method uses images of adult eyes to generate maps with the pixel classifier ilastik (Berg *et al*., 2019), and quantifies red:white surface area ratios with FIJI (Schindelin *et al*., 2012). We previously verified the validity of this technique by comparison to spectrophotometric measurements of pigment extractions (the classic PEV quantification method), and we implement image analysis again here. Ten images were used to train the ilastik pixel classification to distinguish among red pixels, and white pixels. The prediction maps for red, and white were then imported from ilastik into imageJ where a threshold of 0.5 was implemented to remove low-scoring data. From the black and white masks, surface area measurements were made using the ‘oval/elliptical brush selection’ tool and the ‘measure’ tool. We then generated a normalized total surface area by adding the percentages of red and white. The red fraction was calculated as (red%) / (normalized total). A workflow example is presented in **Supp Fig 1**.

For the “Red Fraction of Eye Surface Area” plots, 90.9% of relevant F_1_ eyes were imaged (3376/3716; **Table 1**). Data presented in **Figs 3C,D** include images of every viable fly.

### *Y*-specific PCR

Primers for Exon1 of *kl-3* were designed with Primer3Plus (Untergasser *et al*., 2012). As predicted, BLAST results on Flybase.org showed the 565 bp amplicon to be *Y*-specific. *kl-3_*F: 5’-GGTCGCTACCAGGAAAAGCT, *kl-3_*R: 5’-[GCCGCCAATTGACCAACATT. 30 cycles of PCR (Denature: 95°C for 30 seconds; Anneal: 55°C for 30 seconds; Extension 68°C for 20 seconds) were performed prior to gel electrophoresis.

### Wolbachia status in Drosophila stocks

PCR was regularly used to verify the *Wolbachia* status in *Drosophila* stocks throughout the experiments (Gels in **Supp Fig 5A**) (Cooper & Shropshire, 2024). We used *Wolbachia-*specific PCR primers for *wsp*, and host-specific 28S primers as a positive control. Genomes were extracted in squish buffer, using 3 adult females per reaction. Briefly, flies were homogenized in 50 µL squish buffer per fly (10 mL 1 M Tris-HCl, 0.0372 g EDTA, 0.1461 g NaCl, 90 mL H2O, 150 mL proteinase K), incubated at 65°C for 45 min, incubated at 94°C for 4 min, and centrifuged for 2 min, and the supernatant was used immediately for PCR.

Symbiotic stocks were also verified to be *w*Mel, as opposed to *w*MelCS by PCR amplification of the IS5-WD1310 locus F: 5’-AGGAGAACTGGTCTACGC R: 5’-TGTTGCTGAGCTTTGCT (Gel in **Supp Fig 5B**) (Riegler *et al*., 2005).

### Tetracycline treatment

A 25mM solution was made by dissolving 0.06 grams of Tetracycline HCl in 5mL distilled water. Then, 250µL solution was mixed into 10mL of liquid hot fly food in a food vial (final concentration = 625 µM) and allowed to cool and solidify. The *yw* stock was reared on tet-treated food for three generations, then allowed to recover for two more generations on normal fly food. Finally, *Wolbachia* status was tested by PCR before flies were used in crosses (PCR in **Supp Fig 5A**).

### Figure generation and statistics

Figures were created using Adobe Illustrator (Adobe, San Jose, CA, USA) and in R were used to compare distributions without assuming normalcy. Wilcoxon (Mann-Whitney) tests were two-sided tests. Asterisks on plots mark significance, * *p* <0.05; ** *p* <0.01; *** *p* <0.001; **** *p* <0.0001; ns= not significant or *p* >0.05. Data are presented as box and whisker plots (median ± upper and lower quartiles), overlaid with all the data points for Figures 2C and 5E. PEV comparisons were analyzed using beta generalized linear mixed models (beta GLMM; R v4.3.3, mgcv package, betar() family) with treatment as a fixed effect and experimental cluster (vial or date) as a random intercept, to account for non-independence of flies sharing a vial or experimental day. Red fraction values at exact 0 or 1 were adjusted using the Smithson and Verkuilen (Smithson & Verkuilen, 2006) transformation prior to model fitting. Figures 3C, 3D, and the corresponding subsets of Figure 4A (UU and CI) had vial-level tracking, so vial served as the random effect; for other within-experiment comparisons, experimental date was the clustering variable. Where pairwise comparisons spanned separate experimental blocks with no shared clusters — specifically, comparisons between groups from the *yw* tetracycline-cured experiment (January 2026) and the *yw* symbiotic experiment (September–October 2025) — Mann-Whitney U tests were used, as flies from different experiments are independent. Mann-Whitney was also used where clustering data were unavailable or where experimental date blocks were confounded with treatment (Fig 4B, CI vs Rescue). Multiple pairwise comparisons within each figure were corrected using the Benjamini-Hochberg procedure to control the false discovery rate. Egg-to-adult viability (Table 2) was analyzed using a binomial generalized estimating equation (GEE) with a logit link and exchangeable correlation structure, with experimental day as the clustering factor. Possible correlation between CI strength and PEV suppression was assessed by Pearson correlation on vial-level means (**Supp** Fig 3).

### DAPI staining and imaging

Salivary Glands from third instar larvae were dissected in a depression slide in room temperature PBS. Glands were transferred to a well with mixture of 3:1 ethanol : acetic acid for ∼1 minute fixation, then transferred to a 10 μl drop of 45% acetic acid on a microscope slide. A siliconized coverslip was placed on the drop, and the tissue was lightly squashed between filter paper to draw out the excess acetic acid. The slide was inverted, and flash frozen on dry ice for 10 minutes, coverslip down. Then, a razor blade was used to pry the coverslip off, and the slide was left to dry. A 10 μl drop of Vectashield mixed with 4’,6-diamidino-2-phenylindole (DAPI) to coverslip was placed on top. Once the vectashield had reached the edges of the coverslip, it was sealed to the slide using nail polish. Images were taken on a Zeiss observer 7, using 40x, and 100x objectives. (Protocol adapted from (Kolesnikova *et al*., 2020)). Brain squashes for larval karyotypes were prepared as described by Gatti and Pimpinelli (Gatti & Pimpinelli, 1983).

## Data Availability

All data and code will be made publicly available prior to publication.

## Funding

Research was supported by National Science Foundation (NSF) CAREER (2145195) and National Institutes of Health (NIH) MIRA (R35GM124701) awards to BSC. National Institutes of Health (NIH) MIRA (R35GM139595) to WS. National Institutes of Health (NIH) F32 (F32GM156085) to HJH. Additional support was provided by the University of Montana Genomics Core (UMGC) and Montana INBRE Data Science Core, which are funded by the National Institute of General Medical Sciences (P20GM103474), the Office of the Vice President for Research and Creative Scholarship at the University of Montana, and the M. J. Murdock Charitable Trust (including award 202324717 to BSC). The content is solely the responsibility of the authors and does not necessarily represent the official views of the NSF, UMGC, or the NIH.

## Competing Interests

The authors declare no competing interests.

## Author Contributions

Conceptualization: HJH, WS, BSC; Data curation: HJH; Formal analysis: HJH, BSC; Funding acquisition: WS, BSC; Investigation: HJH; Methodology: HJH,WS, BSC; Project administration: WS, BSC; Resources: WS; BSC; Software: HJH, BSC; Supervision: WS, BSC; Validation: HJH, BSC; Visualization: HJH; Writing – Original draft: HJH; Writing – Review & editing: HJH, WS, BSC.

## Notes

### Competing Interest Statement

The authors have declared no competing interest.

