## Supplemental Figures for "*Wolbachia*-induced cytoplasmic incompatibility produces heritable chromatin modifications that suppress position-effect variegation"

**S1**

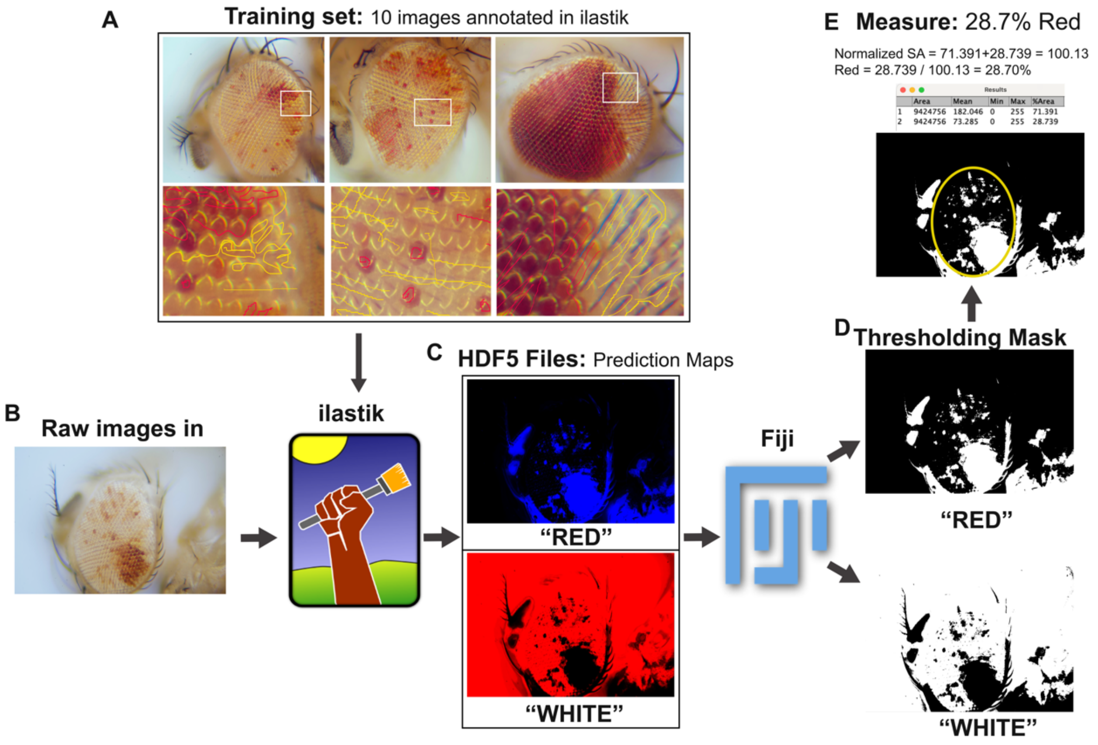


**Supplemental Figure 1:**
**Machine-learning pipeline for quantifying eye pigment from PEV images.** (**A**) Ten representative eye images were used to train the ilastik pixel classifier to distinguish red from white pixels. (**B**) Raw images were imported into ilastik. (**C**) ilastik exported prediction maps (HDF5), which were imported into FIJI (macro A). (**D**) In FIJI, a threshold of 0.5 removed low-confidence pixels (macro B), generating a binary mask for each channel. (**E**) Surface-area measurements were taken from the masks using the oval/elliptical brush selection and measure tools; red fraction was calculated as red area / normalized total area. Macros A (HDF5 import) and B (thresholding and mask conversion) are provided below.

**S1- macros**

**A: Importing multiple HDF5 files to FIJI**

```
// tested with ilastik plugin version 1.8.2

#@ File (label = "Input directory", style = "directory") input_dir

processFolder(input_dir);

// function to scan folder to find files with correct suffix
function processFolder(input_dir) {
	suffix = ".h5";
	list = getFileList(input_dir);
	list = Array.sort(list);
	for (i = 0; i < list.length; i++) {
		if(endsWith(list[i], suffix))
			processFile(input_dir, list[i]);
	}
}

function processFile(input, file) {
	inputFilePath = input + File.separator + file;
	print("Processing: " + inputFilePath);
	run("Import HDF5", "select=[" + inputFilePath + "] datasetname=/exported_data
axisorder=yxc");
}
```

**B: Thresholding HDF5 files and convert to mask**

```
macro "Thresholding [1]" {
setAutoThreshold("Default dark no-reset");
//run("Threshold...");
setThreshold(0.5000, 1000000000000000000000000000000.0000);
setOption("BlackBackground", true);
run("Convert to Mask", "background=Dark calculate black create");
}
```

S2

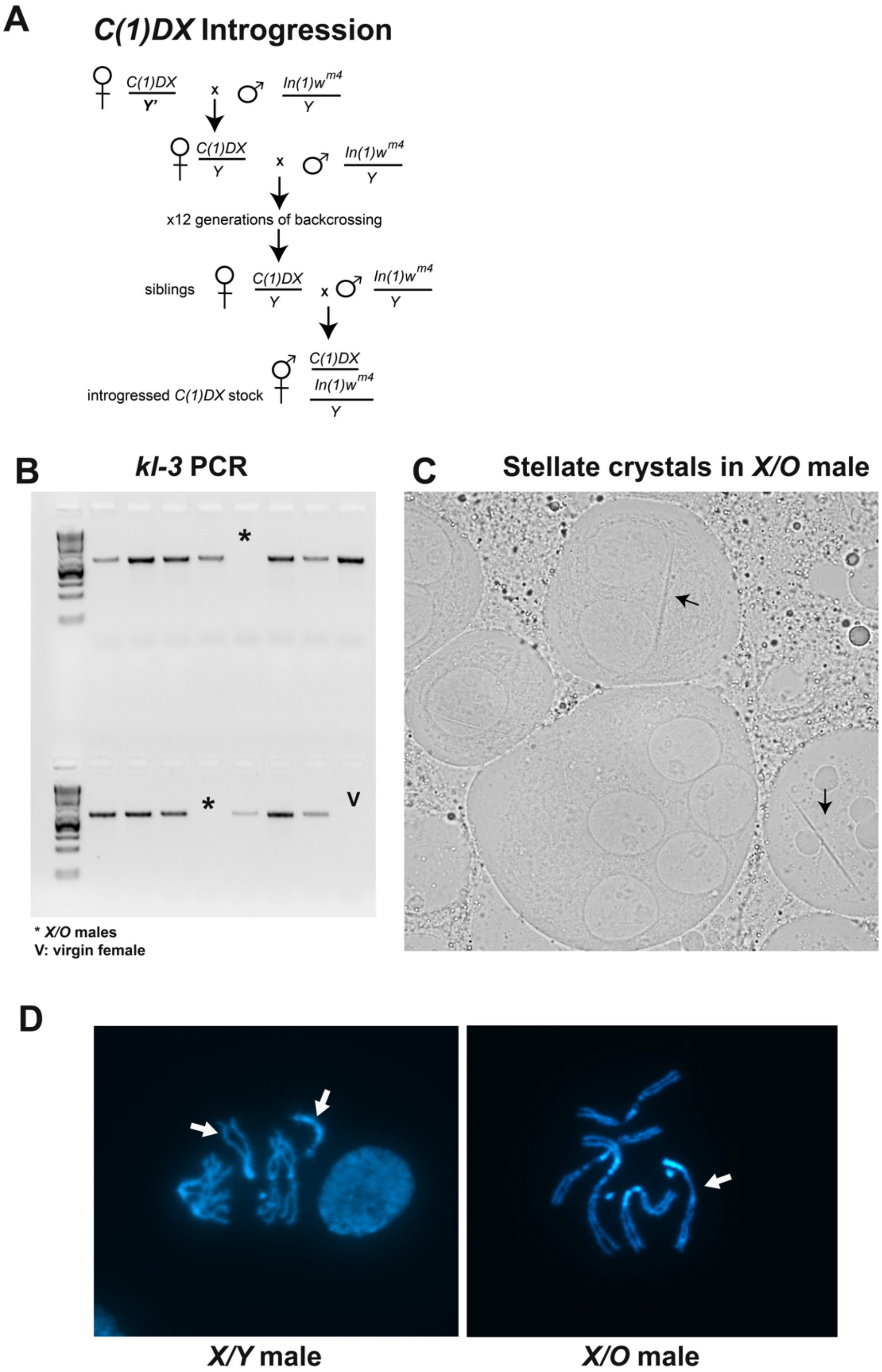


**Supplemental Figure 2 (legend on next page):**

**Supplemental Figure 2:**

**Introgression of the *C(1)DX* background and verification of *X/Y* versus *X/O* patroclinous sons.** (**A**) Crossing scheme for repeated backcrossing of *Wolbachia*-free $In(1)w^{m4}$ males into the *C(1)DX* stock; the *Y* chromosome is introgressed in the first cross. (**B**) Representative PCR genotyping of 15 individual $F_1$ males from a post-introgression CI cross (*C(1)DX* mothers × symbiotic $In(1)w^{m4}$ fathers) using *kl-3* primers targeting the *Y* chromosome. Asterisks mark *Y*-negative lanes, corresponding to *X/O* males; V, virgin-female negative control. (**C**) Brightfield image of a testis from an $F_1$ male derived from a pre-introgression *C(1)DX* cross, hand-selected for severely enhanced variegation. Arrows indicate stellate aggregates (proteinaceous crystals) characteristic of *X/O* males. (**D**) DAPI-stained karyotypes from third-instar larval brains in a *C(1)DX* cross; arrows indicate sex chromosomes. Of 8 male brains karyotyped, 1 was *X/O* and 7 were *X/Y* (12.5%), in line with the proportion of *X/O* males identified by pigment in **Fig 1F** (21/194 = 10.8%).

## S3

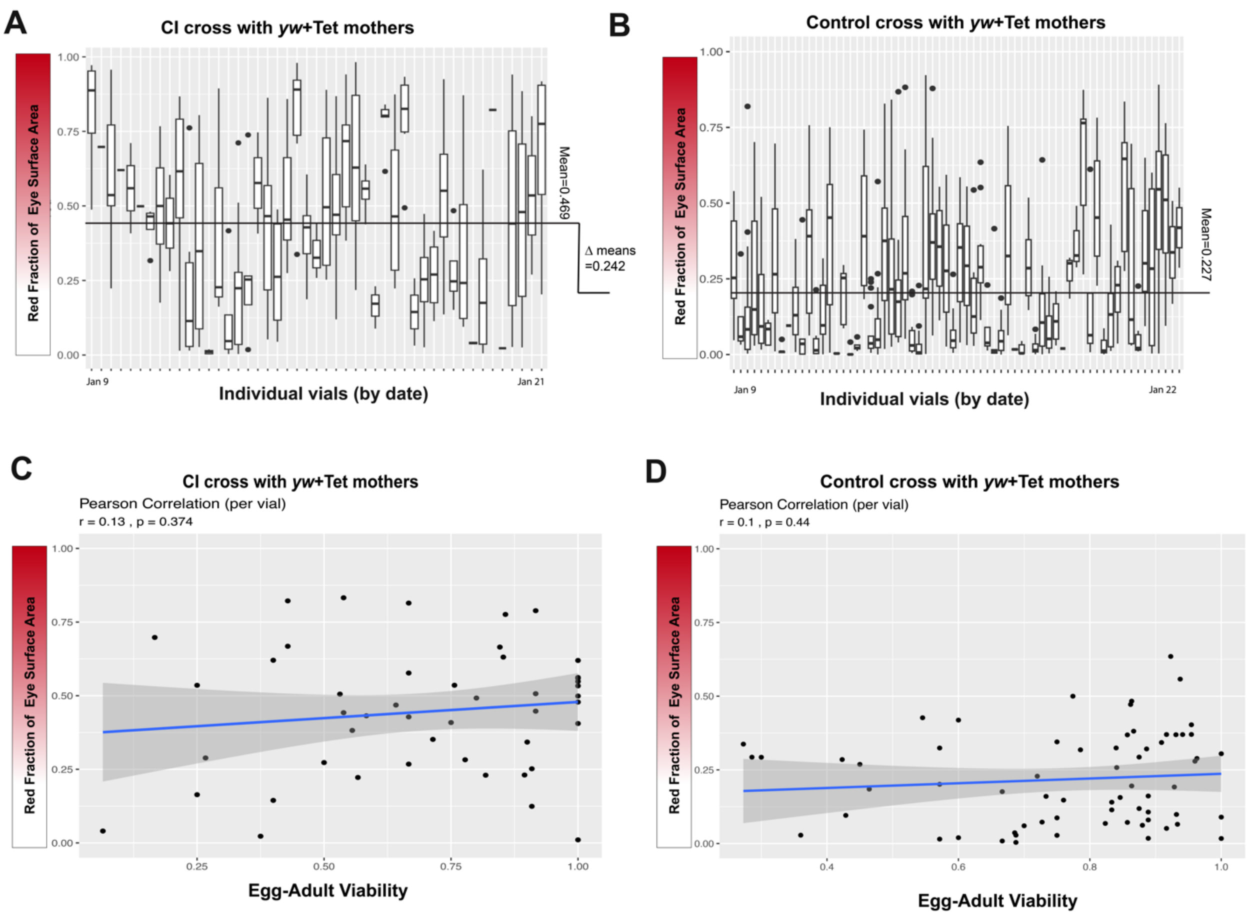


**Supplemental Figure 3:**
**CI strength does not correlate with PEV suppression in the *yw* viability assay.** (**A**) CI-cross vials ordered chronologically; each group of points represents one vial, and the horizontal line marks the overall mean red fraction across all eyes. (**B**) As in (A), for the Control cross. The difference between CI and Control means (0.242) is indicated between the panels. (**C**) Vial-level Pearson correlation between mean eye pigment and egg-to-adult viability for the CI cross ($r$ = 0.13, $p$ = 0.37). (**D**) As in (C), for the Control cross ($r$ = 0.10, $p$ = 0.44). Neither correlation was significant.

**S4**

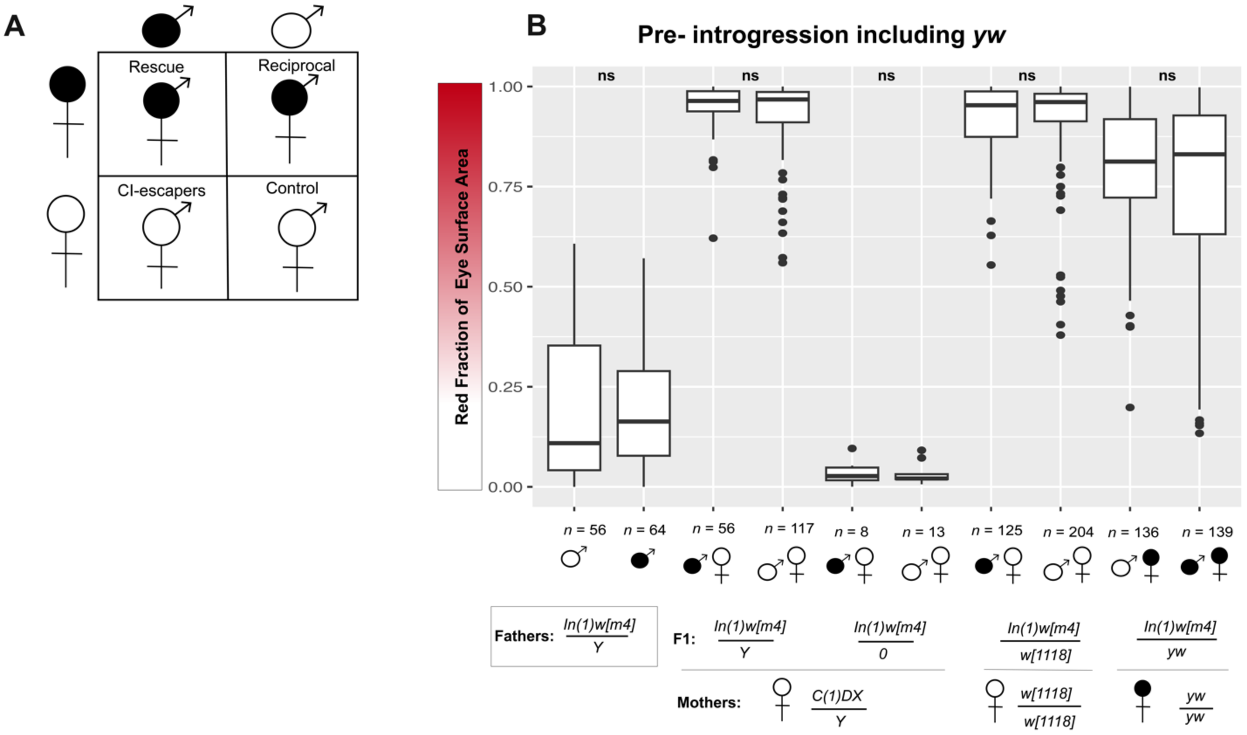


**Supplemental Figure 4:**
**Cross-direction definitions and combined pre-introgression PEV results including symbiotic *yw* crosses.** (**A**) Punnett square defining the four cross directions by parental and offspring *Wolbachia* status, with common names (Control, CI, Reciprocal, Rescue). Filled circles denote *Wolbachia*-carrying individuals, open circles denote *Wolbachia*-free. (**B**) Pre-introgression PEV data from **Fig 1F** with symbiotic *yw* crosses added. Pigment comparisons among $In(1)w^{m}/Y$ males, $In(1)w^{m4}/w^{1118}$ females, and $In(1)w^{m4}/yw$ females used beta GLMMs; comparisons between aposymbiotic and symbiotic fathers and $In(1)w^{m4}/O$ males used Wilcoxon tests. No comparison was significant after BH adjustment ($p_{1\text{-}4} = 0.79$), except the symbiotic *yw* comparison, which was marginal $p_5 = 0.055$. The overall Kruskal-Wallis test across all groups was unchanged from **Fig 1F** ($p < 2.2 \times 10^{-16}$).

**S5**

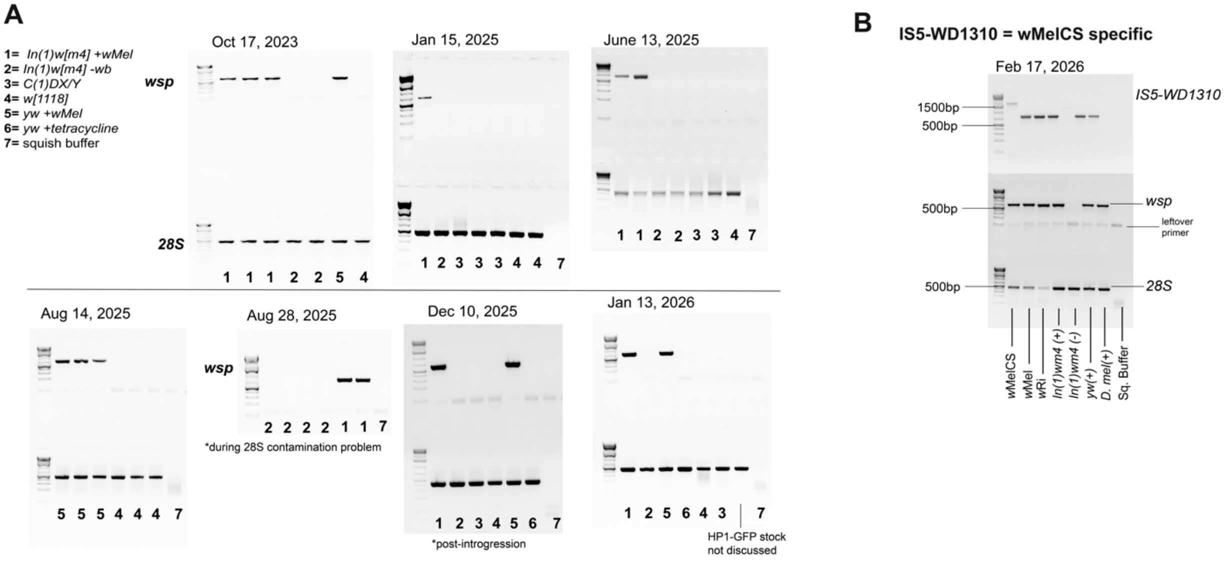


**Supplemental Figure 5:**
**PCR verification of *Wolbachia* infection status and variant identity in *Drosophila* stocks.**
(**A**) Diagnostic PCR for *Wolbachia* infection; numbers in the legend correspond to the genotypes labeled on each dated gel. Template was extracted in squish buffer from 3 adult females per reaction. *wsp* primers are *Wolbachia*-specific (top of each gel); host *28S* primers serve as a positive control for successful extraction (bottom). (**B**) Symbiotic stocks were confirmed as *w*Mel rather than *w*MelCS by PCR amplification of the IS5–WD1310 locus.
